# The TRPC6-AMPK pathway is involved in cytoskeleton reorganization and glucose uptake in podocytes

**DOI:** 10.1101/278911

**Authors:** Patrycja Rachubik, Maria Szrejder, Dorota Rogacka, Irena Audzeyenka, Michał Rychłowski, Stefan Angielski, Agnieszka Piwkowska

**Author notes:** Correspondence author: Agnieszka Piwkowska, Laboratory of Molecular and Cellular Nephrology, Mossakowski Medical Research Center Polish Academy of Sciences, Gdańsk, Poland.

## Abstract

Podocytes are dynamic polarized cells on the surface of glomerular capillaries that are an essential part of the glomerular filtration barrier. AMP-activated protein kinase (AMPK), a key regulator of glucose and fatty acid metabolism, plays a major role in obesity and type 2 diabetes. Accumulating evidence suggests that TRPC6 channels are crucial mediators of calcium transport in podocytes and are involved in regulating glomerular filtration barrier. Here we investigated whether the AMPK-TRPC6 pathway is involved in insulin-dependent cytoskeleton reorganization and glucose uptake in cultured rat podocytes. Insulin regulates the interaction of TRPC6 with AMPKα2 in cultured rat podocytes The results suggested a key role for the TRPC6 channel in the mediation of insulin-dependent activation of AMPKα2, actin cytoskeleton reorganization and glucose uptake in podocyte. Moreover, AMPK and TRPC6 activation were required to stimulate the Rac1 signaling pathway. These results suggest a potentially important new mechanism that regulates glucose transport in podocytes and that could be injurious during diabetes.

## Introduction

The impairment of insulin signaling and insulin pathways in skeletal muscle, fat tissue, and the liver is central to the development of type 2 diabetes. The resulting dysregulation of glucose and fat metabolism leads to kidney failure and/or to cardiovascular complications. Indeed, the majority of patients with albuminuria and end-stage renal failure in Western countries show abnormalities in insulin production or in insulin effectiveness.(Rawshani & Gudbjornsdottir, 2017, Ritz, Rychlik et al., 1999

Podocytes are uniquely sensitive to insulin and have similarities to skeletal muscle and fat cells with respect to insulin-stimulated glucose uptake kinetics and the expression of glucose transporters (GLUTs) (Coward, Welsh et al., 2005, Rogacka, Piwkowska et al., 2016). Recent studies have shown that insulin can dynamically remodel the actin cytoskeleton of podocytes and that this is critically important for maintaining the integrity of the glomerular filtration barrier. Actin reorganization leads to changes in podocyte structure, and insulin receptor stimulation causes the retraction of podocyte processes (Welsh, Hale et al., 2010). Some groups have suggested that insulin plays a role in the control of podocyte contractility, which may contribute to glomerular permeability (Kim, Anderson et al., 2012, Kim & Dryer, 2011, Piwkowska, Rogacka et al., 2015, Piwkowska, Rogacka et al., 2013). We recently demonstrated that insulin remodels the actin cytoskeleton and increases the albumin permeability of both isolated rat glomeruli and podocytes; we further showed that the underlying mechanism is calcium-dependent (Piwkowska, Rogacka et al., 2016, Rogacka, Audzeyenka et al., 2017). Moreover, in mouse podocytes, insulin increases the steady-state cell surface expression of TRPC6 channels (Kim et al., 2012).

Overexpression or gain of function of the TRPC6 channel can drive podocyte effacement, and TRPC6 overexpression in the mouse kidney induces proteinuria (Krall, Canales et al., 2010). An increase in intracellular calcium downregulates the expression of nephrin and synaptopodin and stimulates RhoA activity, which in turn causes F-actin derangement and a decrease in foot processes (Jiang, Ding et al., 2011). Furthermore, TRPC6 gene mutations are linked to human proteinuric kidney disease, focal segmental glomerulosclerosis, and loss of podocytes (Dryer & Reiser, 2010).

We postulated that dynamic responses in podocytes and in their foot processes are mediated by cytoskeletal elements and by Ca^2+^-dependent processes. However, the means by which proteins regulate actin dynamics in podocyte foot processes are not fully understood, especially in diabetes. In many types of glomerular diseases, the integrity of the actin cytoskeleton is altered in podocytes, indicating that proper organization and regulation of the actin cytoskeleton are essential for podocyte structure and function (Mathieson, 2011).

AMP-activated protein kinase (AMPK) activity appears to positively regulate insulin-dependent glucose uptake and insulin signaling (Fisher, Gao et al., 2002, Jing, Cheruvu et al., 2008); however, AMPK activity is suppressed in disorders associated with insulin resistance (Rogacka, Piwkowska et al., 2014, Rogacka et al., 2016). The AMPK is composed of three subunits, one catalytic, termed α, and two regulatory, termed β and γ. The activation of AMPK requires the phosphorylation of threonine 172 (Thr172) within the catalytic α subunit by upstream kinases, namely the Ca^2+^/calmodulin-dependent kinase kinase-β (CaMKK-β) and/or the LKB1–STRAD–MO25 complex (Zeqiraj, Filippi et al., 2009). Hypoxia and contractile activity also activate AMPK. In addition to the potential requirement for AMPK activity, normal regulation of contraction- and exercise-stimulated glucose uptake also requires the Rho GTPase Rac1 (Sylow, Moller et al., 2015, Sylow, Moller et al., 2017), which is activated by insulin and which induces actin cytoskeleton remodeling at the plasma membrane in skeletal muscle cells (Sylow, Jensen et al., 2013). Rac1 mediates this process by inducing cortical F-actin remodeling, which involves the recruitment of actin regulatory proteins such as cofilin and Arp2/3 to the actin filaments (Chiu, Patel et al., 2010). In addition, Rac1 signals p21-activated kinase 1 (PAK1) in skeletal muscle and facilitates PAK1 phosphorylation in response to insulin (Tsakiridis, Taha et al., 1996). Insulin-stimulated PAK1 activation is decreased in human skeletal muscle in both acute (intralipid infusion) and chronic (obesity and type 2 diabetes) insulin resistant states (Sylow et al., 2013), suggesting that PAK1 is a required element for maintaining euglycemia and insulin sensitivity. These findings suggest that Rac1 and downstream signaling to the actin cytoskeleton constitute an important dysfunctional pathway in insulin-resistant states. This mechanism not be recognized in podocytes.

In the present study, we investigated whether the TRPC6-AMPK pathway is involved in insulin-mediated cytoskeleton reorganization and glucose uptake. Our results identified a potentially important new mechanism that may be injurious to podocytes in diabetes and consequently interfere with the intracellular transport of glucose.

## Results

### Intracellular calcium signaling regulates insulin signal transduction in cultured rat podocytes

First, we investigated the effects of intracellular calcium signaling on the phosphorylation of proteins involved in insulin signal transduction in cultured rat podocytes (Fig. 1). Insulin treatment (300 nM, 5 min) increased the phosphorylation of the insulin receptor (IR) by 262% (from 0.34±0.07 to 1.23±0.24, n=4, *P*<0.05) and increased the phosphorylation of Akt by 32% (from 1.69±0.04 to 2.24±0.03, n=4, *P*<0.05). Incubation of the podocytes with inhibitors of calcium extrusion, namely La^3+^ (1 mM), caloxin2A1 (CX, 0.3 mM), or CB-DMB (50 μM) had no effect on IR and Akt phosphorylation. Insulin treatment of podocytes in the presence of all of the inhibitors of calcium extrusion increased the phosphorylation of IR and Akt to similar extents.

**Figure 1.**
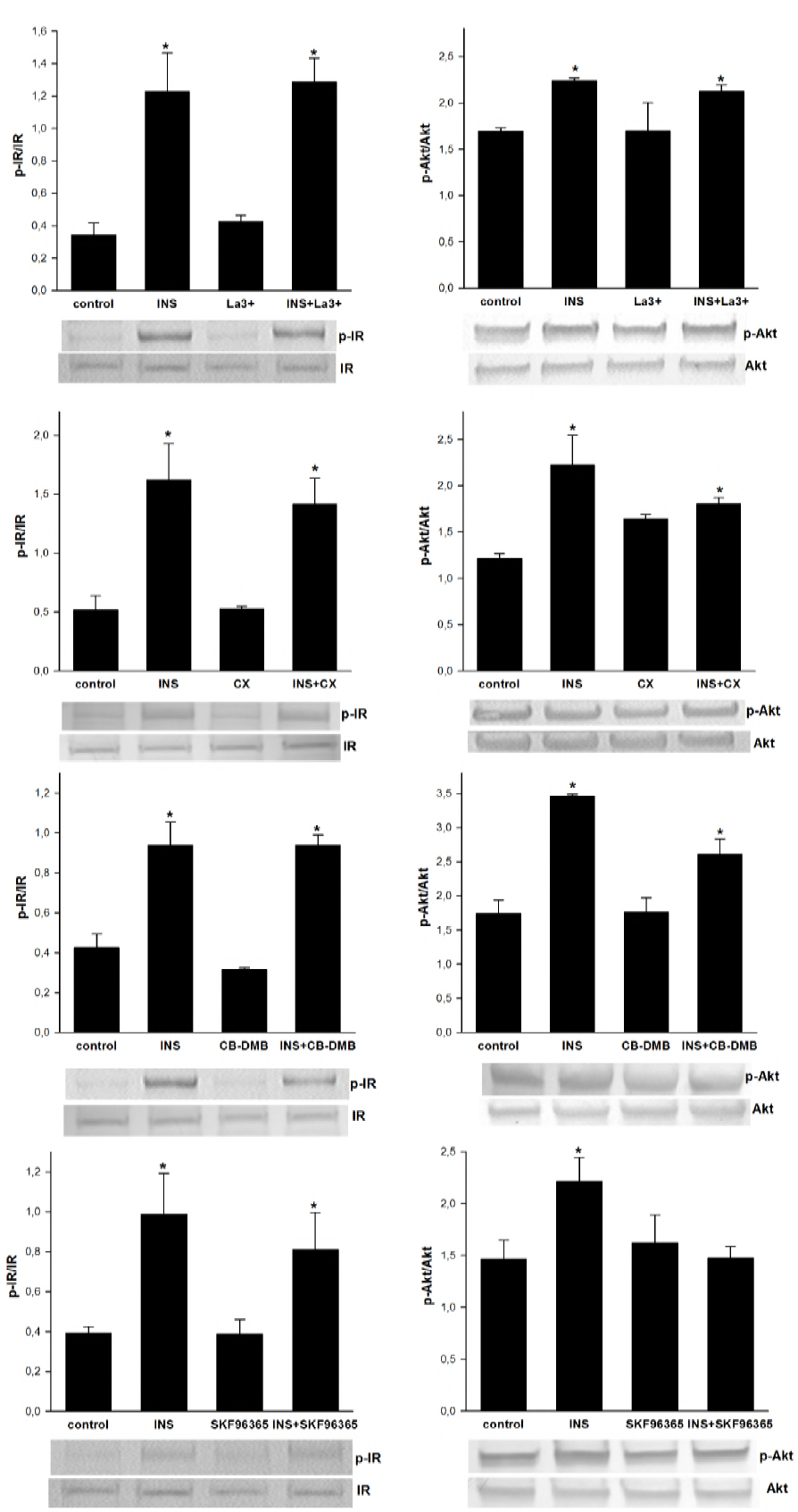
The effect of intracellular calcium signaling on the phosphorylation of proteins involved in insulin signal transduction in cultured rat podocytes. Podocytes were incubated with or without insulin (300 nM, 5 min) and the indicated calcium signaling inhibitors. Cell lysates were analyzed by immunoblotting using anti-IR, anti-p-IR (Tyr^1150/1151^), anti-Akt, and anti-p-Akt (Ser^473^) antibodies. Values are reported as the means±SEMs of four independent experiments. \**P*<0.05 compared to control.

Next, we evaluated the effect of a TRPC inhibitor (SKF96365, 100 μM, 15 min preincubation) on insulin signal transduction in the cultured podocytes. SKF96365 did not affect IR phosphorylation, but it blocked the effect of insulin on Akt phosphorylation (1.47±0.11 vs. control 1.46±0.18, n=4). We then evaluated the role of TRPC6 on insulin signal transduction in podocytes by knocking down TRPC6 expression using siRNA (Fig. 2A). There was a significant decrease in TRRP6 protein expression in podocytes transfected with TRPC6 siRNA compared to podocytes transfected with the control scrambled siRNA (55% decrease, 0.221±0.039 vs. control 0.490±0.013, n=4, *P*<0.05). Downregulation of TRPC6 expression abolished insulin-induced Akt phosphorylation without affecting IR phosphorylation (Fig. 2B, 2C).

**Figure 2.**
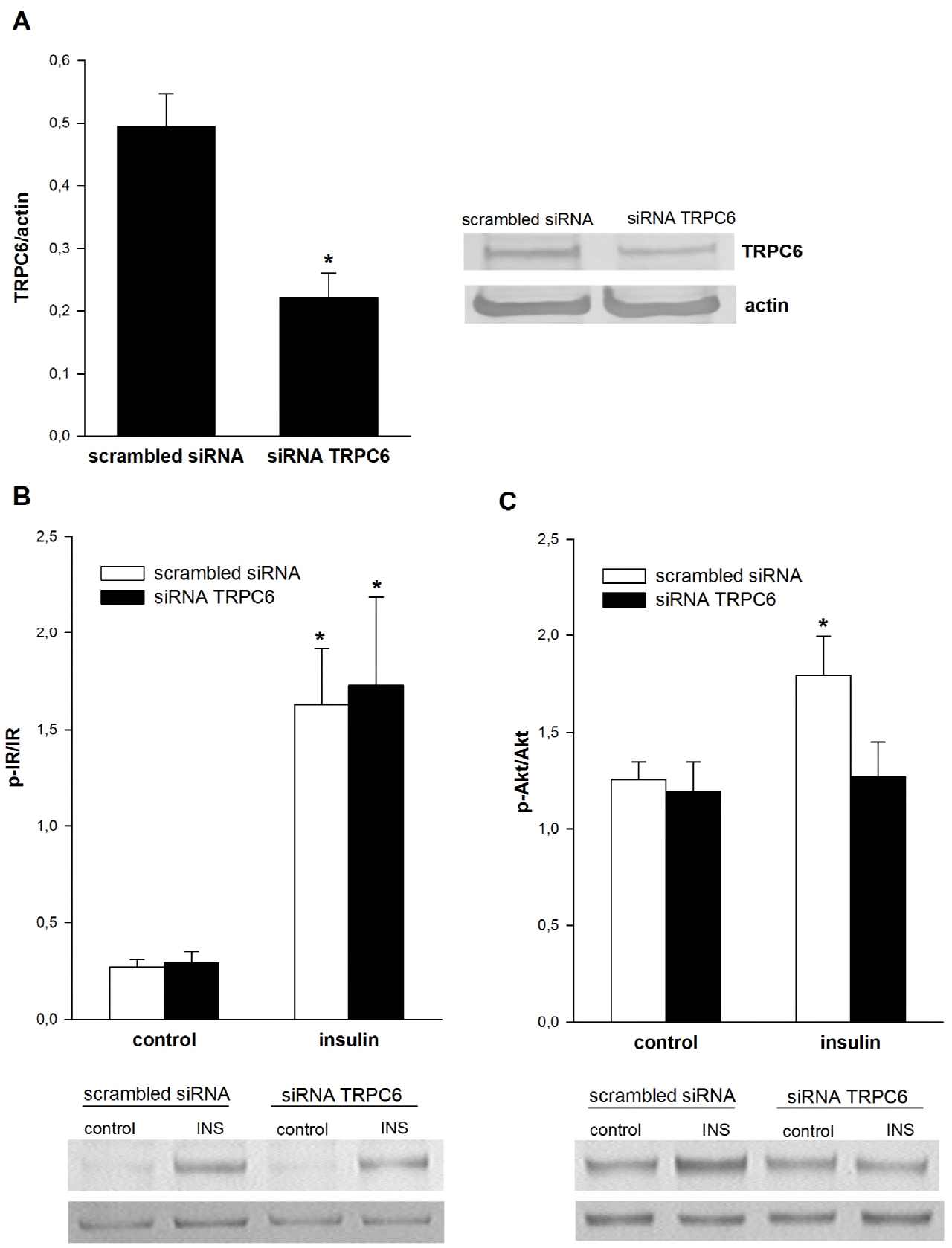
The influence of TRPC6 channels on insulin signal transduction in cultured rat podocytes. (A)The effect of TRPC6 siRNA on TRPC6 expression. Densitometric measurements of the TRPC6 bands were normalized to the actin band. The influence of TRPC6 downregulation on insulin-dependent phosphorylation of insulin receptor (B) and kinase Akt (C). Values are reported as the means±SEMs of four to six independent experiments. \**P*<0.05 compared to appropriate control.

### Intracellular calcium signaling regulates the insulin-dependent activation of AMPK kinase and glucose uptake in cultured rat podocytes

AMP-activated protein kinase (AMPK) is a major regulator of insulin-dependent glucose uptake and insulin signaling in podocytes (Rogacka et al., 2016). We found that insulin (300 nM, 5 min) induced the phosphorylation of the AMPKα subunit (0.799±0.081 vs. control 0.569±0.042, n=4, *P*<0.05, Fig. 3A). We hypothesized that insulin might increase AMPKα phosphorylation by activating TRPC channels. Indeed, preincubation of podocytes with the TRPC channel inhibitor SKF96365 (100 μM, 15 min preincubation) abolished the effect of insulin on AMPK phosphorylation (Fig. 3A). The same effect was observed after downregulation of TRPC6 expression in podocytes (Fig. 3B). We then demonstrated that podocytes were insulin-sensitive and that insulin stimulation increased glucose uptake by about 40% (Fig. 4). To determine the effect of TRPC channels on insulin-dependent glucose uptake, we evaluated the effect of SKF96365 (Fig. 4A) and downregulation of TRPC6 expression (Fig. 4B) and found that both blocked insulin-dependent glucose uptake. We therefore postulated that TRPC6 channels play an essential role in the regulation of insulin-dependent signaling and thus in the regulation of glucose uptake in podocytes.

**Figure 3.**
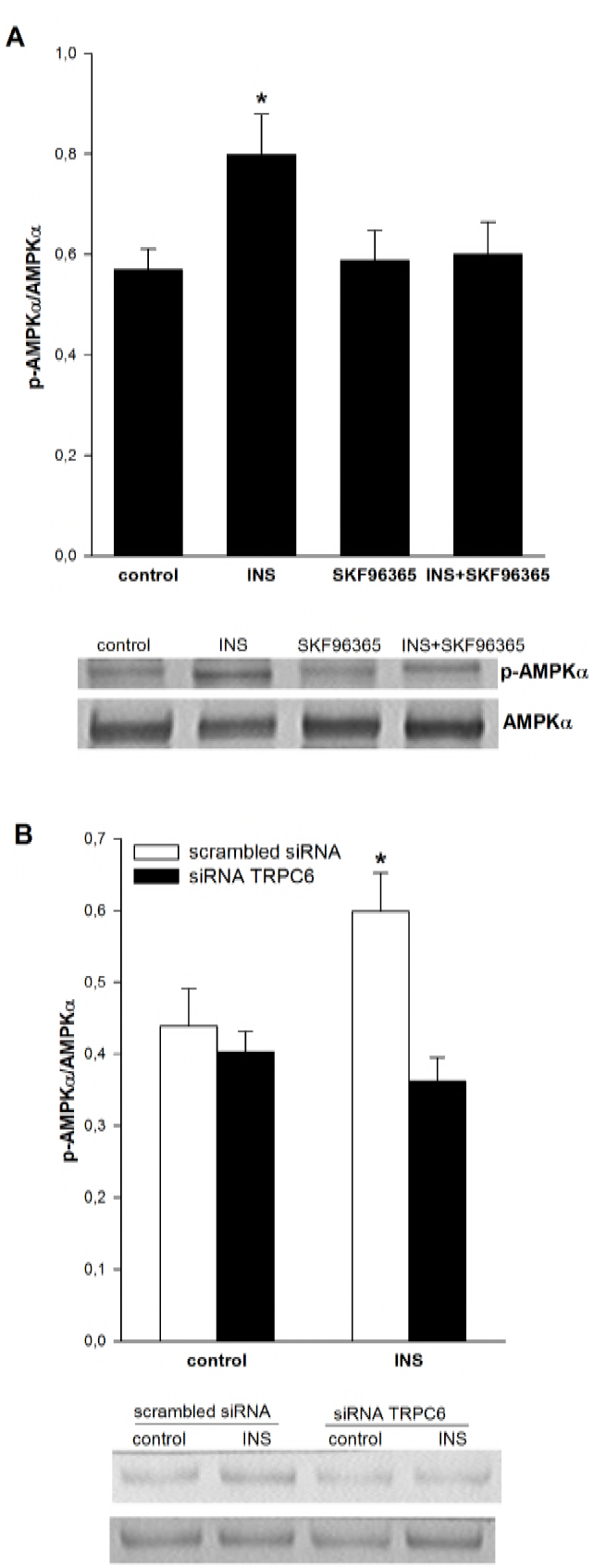
TRPC6 channels mediate insulin-dependent activation of AMPK in cultured rat podocytes. The effects of the TRPC6 inhibitor SKF96365 (A) and of downregulation of TRPC6 channels(B) on the insulin-dependent activation of AMPK. Cell lysates were subjected to immunoblotting analysis using anti-AMPKα and anti-p-AMPKα (Thr^172^) antibodies. Values are reported as the means±SEMs of four independent experiments. \**P*<0.05 compared to control.

**Figure 4.**
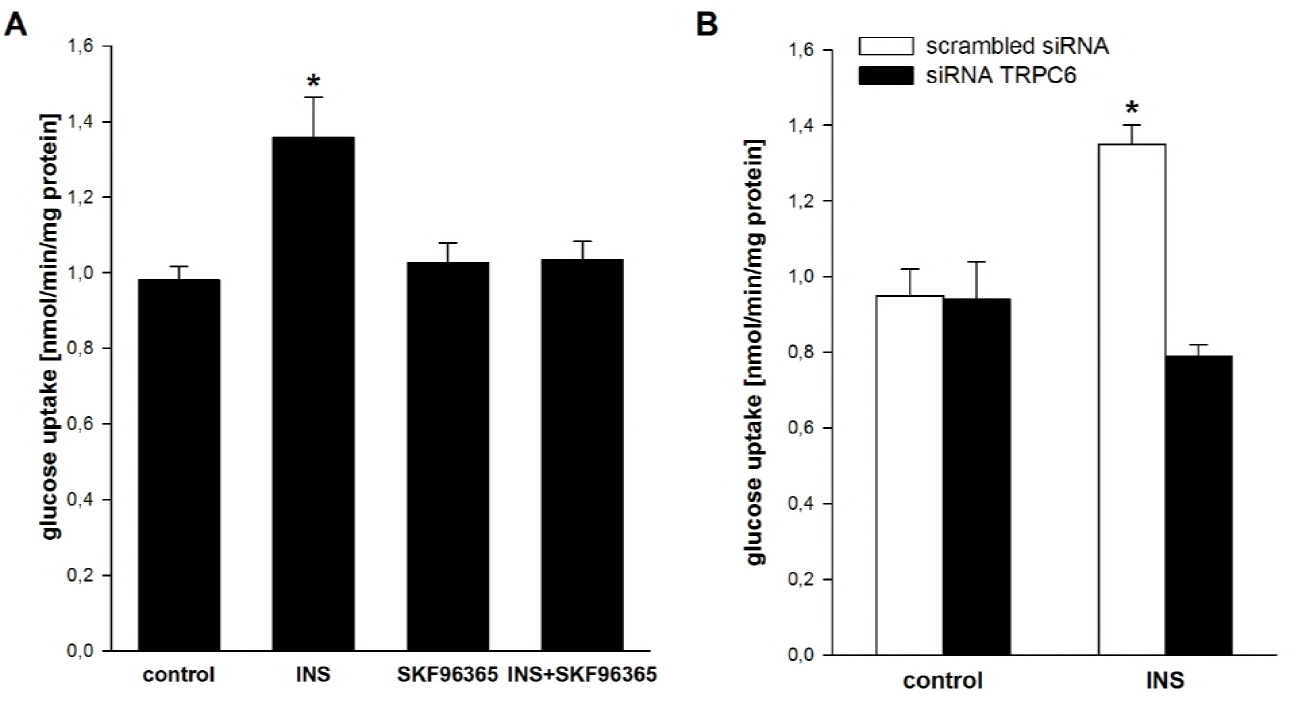
The effect of TRPC6 channels on insulin-dependent glucose uptake in cultured rat podocytes. The effects of the TRPC6 inhibitor SKF96365 (A) and of downregulation of TRPC6 channels (B) on insulin-dependent increases in glucose uptake. Uptake measurements were performed after the addition of 1 μCi of [1,2-^3^H]-deoxy-D-glucose diluted in non-radioactive glucose to a final concentration of 300 nM insulin. Values are reported as the means±SEMs of four to five independent experiments. \**P*<0.05 compared to control.

### The role of insulin in TRPC6 channel interactions with AMPKα subunits in cultured rat podocytes

Many groups have shown that TRPC6 is part of a larger signaling complex in the slit diaphragm and that this complex plays an important role in the regulation of podocyte function. We investigated whether AMPKα subunits associate with TRPC6 as well as the possible role of insulin in this interaction. After mixing podocyte extract with antibodies against AMPKα1 or AMPKα2 subunits, TRPC6 was detected in both immunoprecipitates. Similarly, antibodies to TRPC6 coimmunoprecipitated AMPKα1 and AMPKα2 (Fig. 5A). We also found that insulin increased the amount of TRPC6 that co-immunoprecipitated with AMPKα2 subunits by approximately 45% (from 0.579±0.064 to 0.838±0.055, n=3, *P*<0.05, Fig. 5B). The quantitative analysis confirmed that insulin increased the colocalization of TRPC6 with AMPKα2 subunits (from 57.5% to 72%, n=8, *P*<0.05, Fig. 6). Insulin did not increase the level of TRPC6 that colocalized with AMPKα1 subunits.

**Figure 5.**
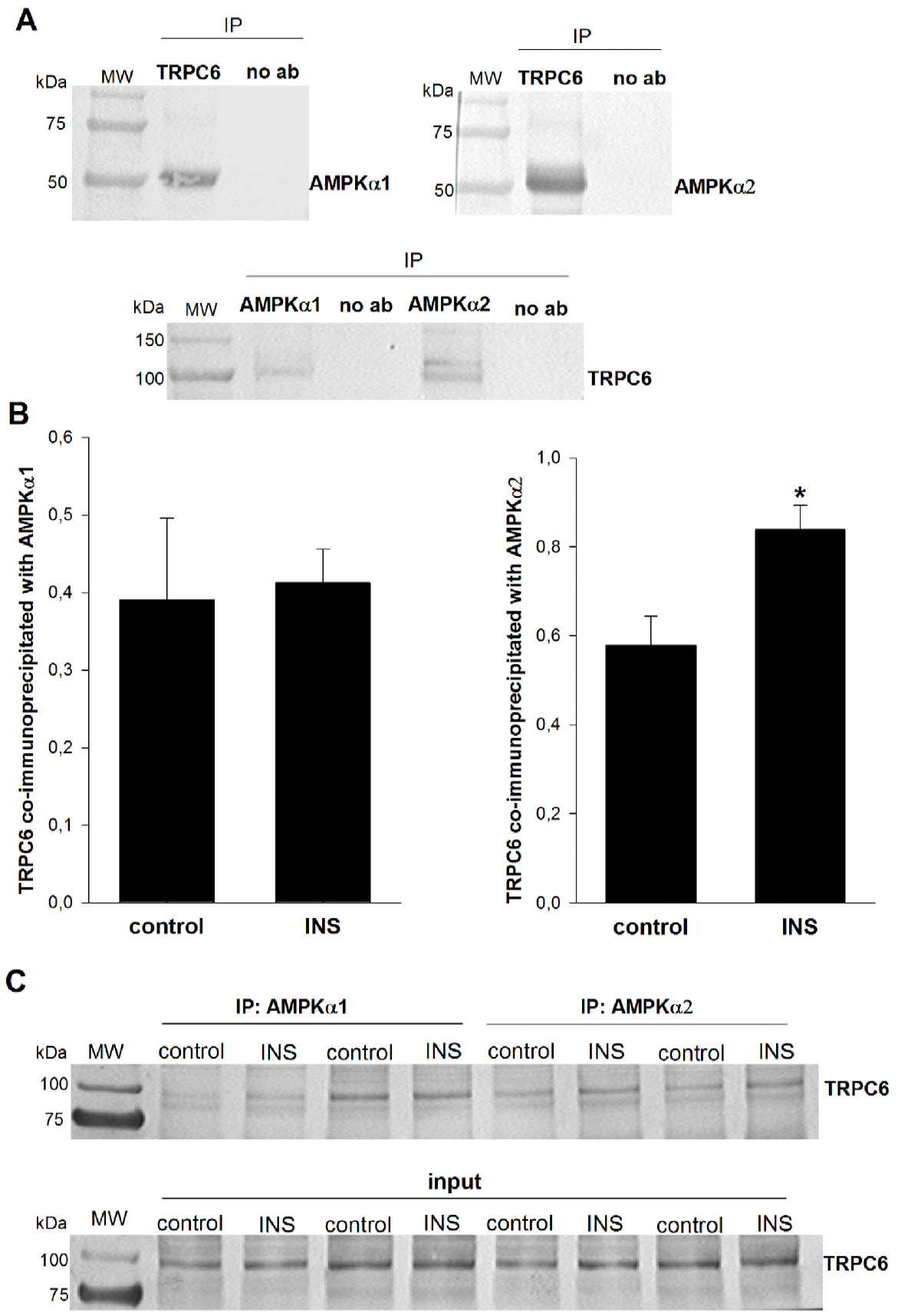
The role of insulin in TRPC6 channel interactions with AMPKα subunits in cultured rat podocytes. (A) Immunoblots showing that the AMPKα1 and AMPKα2 subunits are associated with immunoprecipitated TRPC6 in podocyte extracts and that, conversely, TRPC6 is associated with immunoprecipitated AMPKα1 and AMPKα2. (B) Insulin increases the amount of TRPC6 that coimmunoprecipitates with AMPKα2. (C) Representative immunoblots.

**Figure 6.**
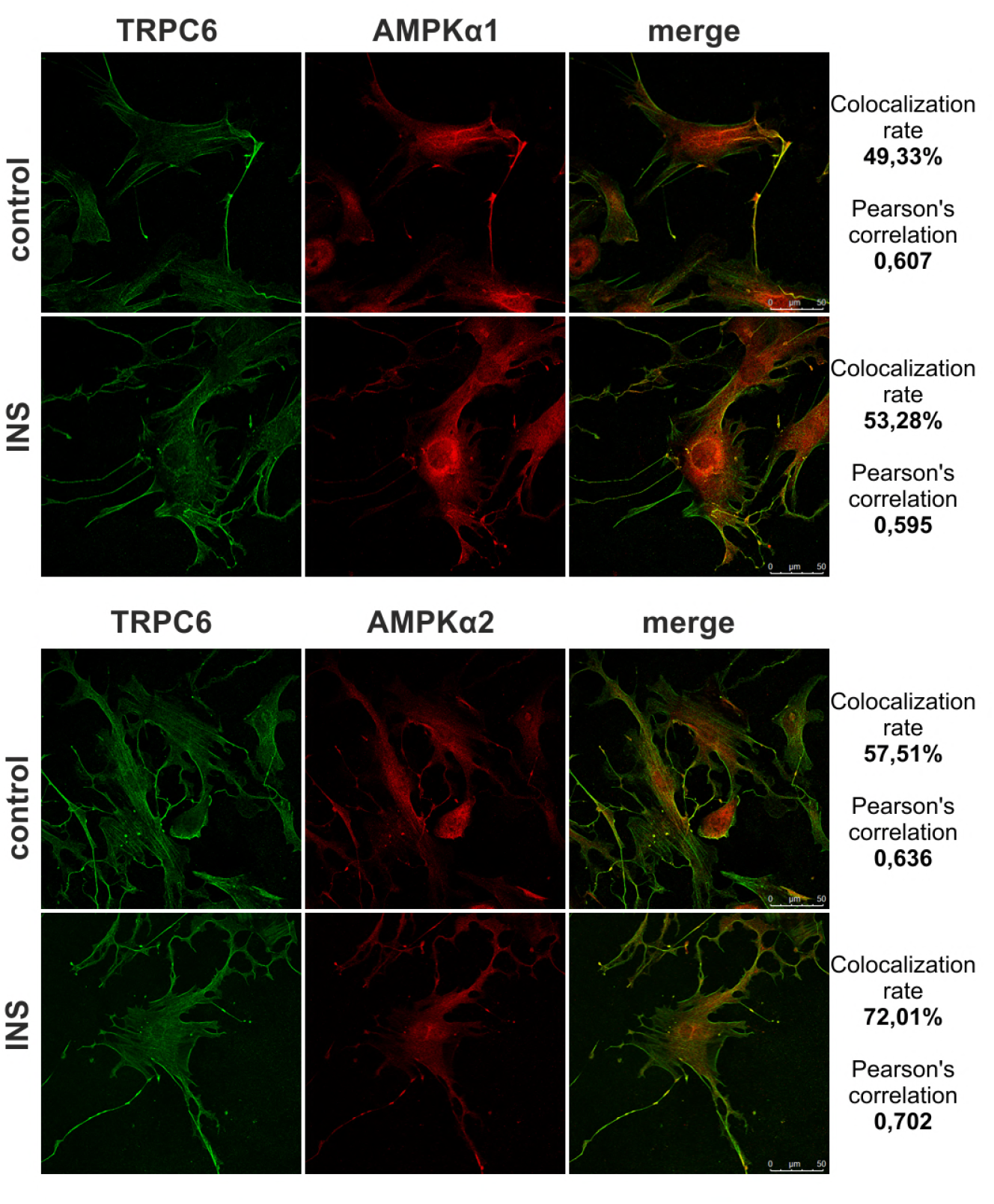
Insulin changes the colocalization of TRPC6 and AMPKα in cultured rat podocytes. Podocytes were seeded onto coverslips and incubated with or without insulin (300 nM, 5 min). The cells were than immunoblotted with anti-TRPC6 and anti-AMPKα1 or anti- AMPKα2 antibodies as indicated. Quantitative analysis of protein colocalization was performed with LAS AF 3.3.0 software (n=8–10). The pixel intensities were quantified and the results are reported as Pearson’s correlation coefficients and colocalization rates (%).

### The role of TRPC6 and AMPK in insulin-dependent regulation of Rac1 activity in podocytes

The small GTPase Rac1 is a major regulator of actin remodeling, and cytoskeletal rearrangement is required for GLUT4 translocation in response to insulin (JeBailey, Wanono et al., 2007). Because we found that inhibition of TRPC6 decreased insulin-stimulated glucose uptake, we hypothesized that this might be due to impaired Rac1-dependent regulation of the actin cytoskeleton. Rac1 is activated when bound to GTP. We found that insulin induces activation of Rac1 by 36% compared to control (1.34±0.02 vs. control 0.86±0.02, n=4, *P*<0.05). Moreover, preincubation of podocytes with TRPC inhibitor (SKF96365, 100 μM, 15 min) attenuated the effect of insulin on Rac1 activity (Fig. 7). To determine the influence of AMPK on Rac1, we modified AMPK kinase activity using the AMPK activator metformin (2 mM) and the AMPK inhibitor compound C (100 μM) (Fig. 8A). We found that metformin increased glucose uptake by 37% (0.993±0.063 vs. control 0.725±0.048, *P*<0.05, Fig. 8B). However, the AMPK inhibitor compound C decreased glucose uptake to 0.617±0.029 (*P*<0.05, Fig. 8B). Similar to insulin, metformin also increased Rac1 activity (Fig. 8C) by 39% (1.21±0.14 vs. control 0.87±0.06, n=6, *P*<0.05, Fig. 8C) and Rac1 serine 71 phosphorylation by 25% (1.53±0.07 vs. control 1.22±0.09, n=5, *P*<0.05, Fig. 8D). Inhibiting AMPK activity with compound C decreased Rac1-GTP binding by 29% and Rac1 phosphorylation by 20%.

**Figure 7.**
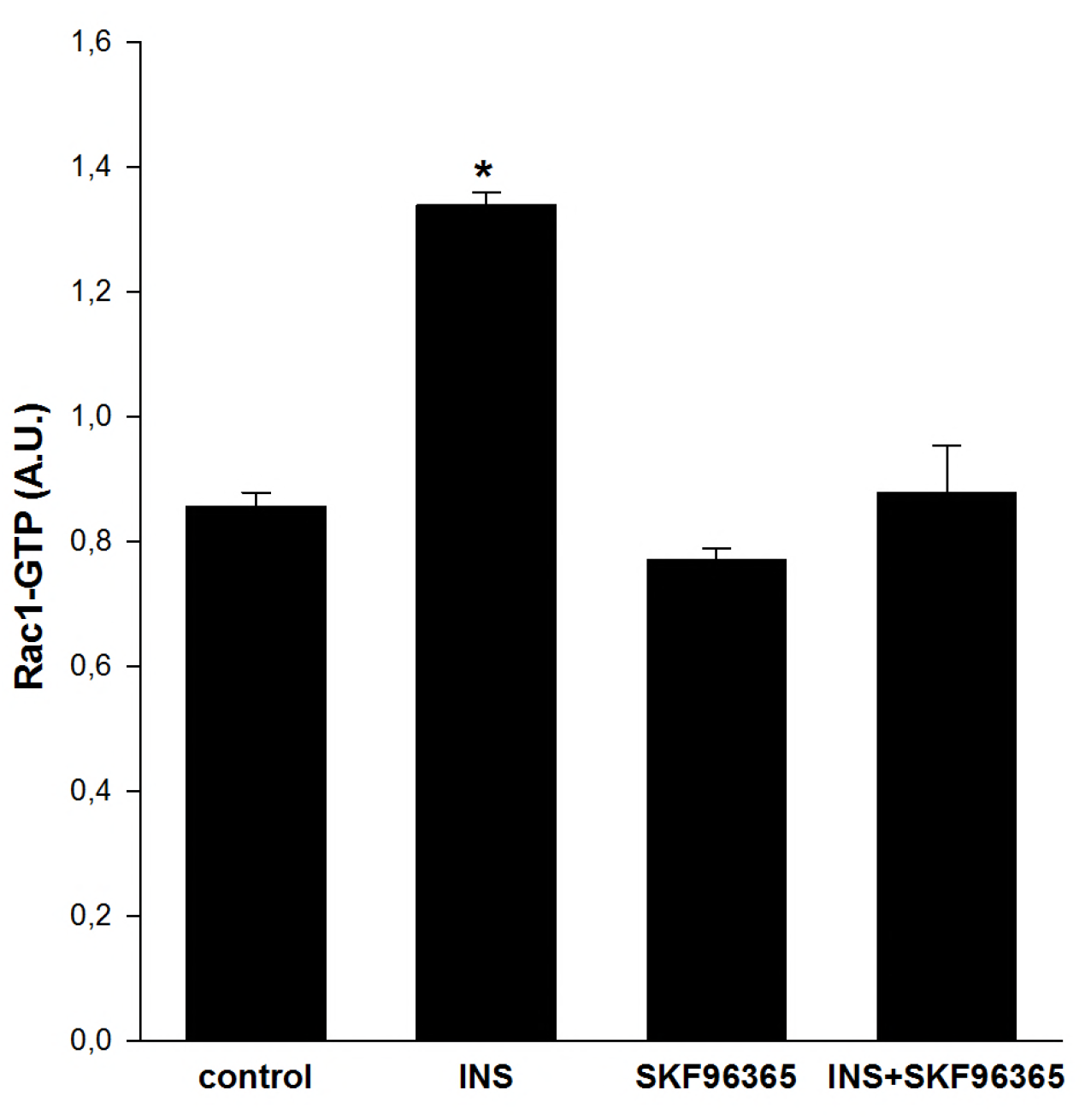
The influence of TRPC channels on Rac1 activity in cultured rat podocytes. Cells were incubated for 5 min with 300 nM insulin in the presence or absence of the TRPC6 inhibitor SKF96365 (10 μM, 20 min). Cell lysates (20 μg) were subjected to the G-LISA assay, and absorbance was read at 490 nm. The data are background-subtracted. Values are reported as the means±SEMs of six independent experiments. \**P*<0.05 compared to control.

**Figure 8.**
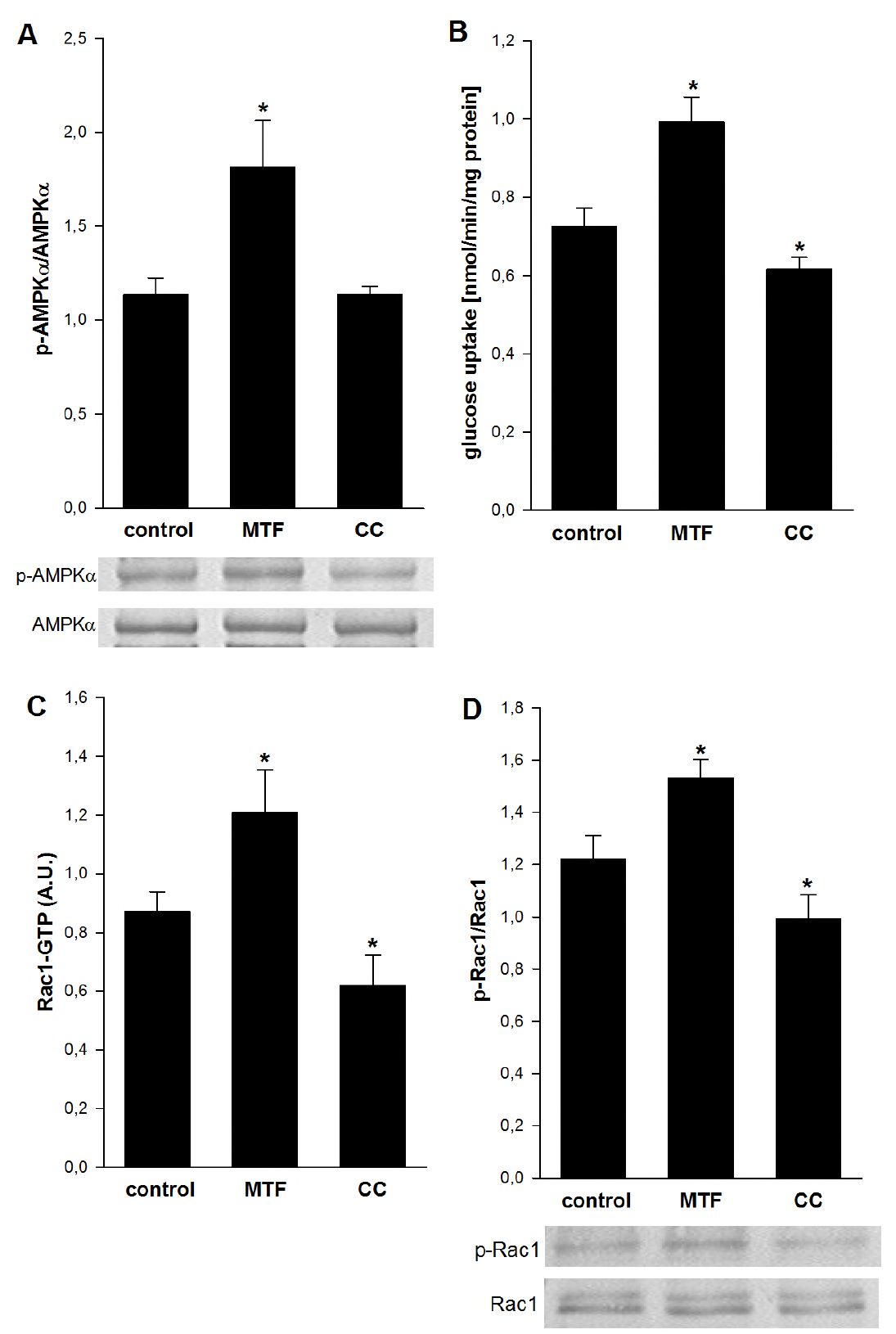
The influence of AMPK on the activity of the small GTPase Rac1 in cultured rat podocytes. The effect of the AMPK activator metformin (2 mM) and the AMPK inhibitor compound C (100 μM) on AMPK activity (A) and on glucose uptake (B). The effect of AMPK on the regulation of Rac1 activity (C) and on serine 71 phosphorylation of Rac1 (D). Values are reported as the means±SEMs of four to six independent experiments. \**P*<0.05 compared to control.

We then evaluated the roles of the AMPKα1 and AMPKα2 subunits in the regulation of Rac1 activity. We knocked down AMPKα1 and AMPKα2 expression using siRNA (Fig. 9A, B).

**Figure 9.**
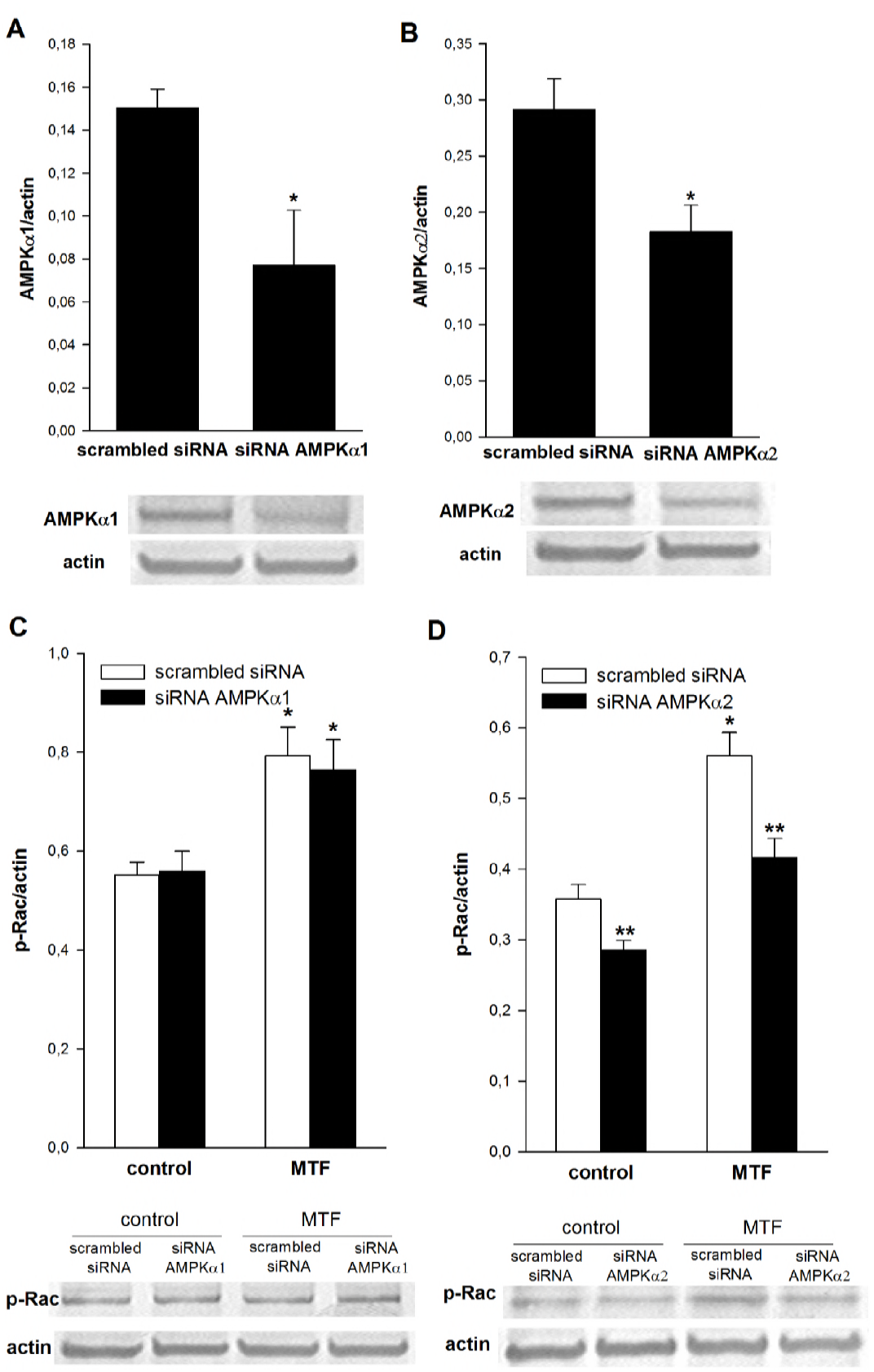
The role of AMPKα1 and AMPKα2 subunits in the regulation of Rac1 activity. The effects of small interfering RNA targeting transcripts of the AMPKα1 or AMPKα2 subunit in cultured rat podocytes. Densitometry was performed to evaluate the expression of AMPKα1 (A) and AMPKα2 (B), and the band signals were normalized using signals of actin bands. Controls were transfected with scrambled siRNA. The influence of downregulation of AMPKα1 (C) and AMPKα2 (D) on Rac1 phosphorylation. Values are reported as the means±SEMs of four independent experiments. \**P*<0.05 compared to control, \*\**P*<0.05 compared to the appropriate control with scrambled siRNA.

Only downregulation of AMPKα2 expression decreased Rac1 phosphorylation in the control (by 21%, *P*<0.05, Fig. 9D) and in the presence of metformin (25%, *P*<0.05, Fig. 9D). These data suggest that activation of both TRPC6 and AMPKα2 are necessary to activate Rac1 in cultured rat podocytes.

### Insulin-mediated activation of downstream targets of Rho kinases is TRPC6-dependent

We next evaluated the influence of TRPC6 on the insulin-dependent regulation of downstream targets of Rho kinases. Insulin-stimulated activation of Rac1 activates the serine/threonine kinase PAK in skeletal muscle (Sylow et al., 2013). Activation of PAK involves phosphorylation of the Thr423 and Ser141 residues, dissociation of the dimer, and release of the catalytic domains (Lei, Lu et al., 2000). In podocytes, insulin stimulation (300 nM, 5 min) increased PAK Ser141 phosphorylation by 71% (1.20±0.24 vs. control 0.70±0.03) and PAK Thr423 phosphorylation by 33% (1.54±0.11 vs. control 1.16±0.09)(Fig. 10, *P*<0.05 for both). Furthermore, we found that preincubating cells with SKF96365 (100 μM) or downregulation of TRPC6 by siRNA attenuated the effect of insulin on PAK phosphorylation in podocytes (Fig. 10). We also investigated the influence of TRPC6 activity on ROCK1 and ROCK2 levels in rat podocytes. ROCK is a major downstream effector of the RhoA kinase (Burridge & Wennerberg, 2004). The TRPC6 inhibitor SKF96365 did not influence ROCK kinase levels (Fig. 11A, C). Moreover, downregulation of TRPC6 by siRNA decreased the ROCK1 level by 23% (from 0.644±0.036 to 0.488±0.014, *P*<0.05, Fig. 11B) without affecting ROCK2 expression. Taken together, these results suggest that downregulation of TRPC6 attenuated the insulin-dependent activation of downstream targets of Rho kinases in podocytes.

**Figure 10.**
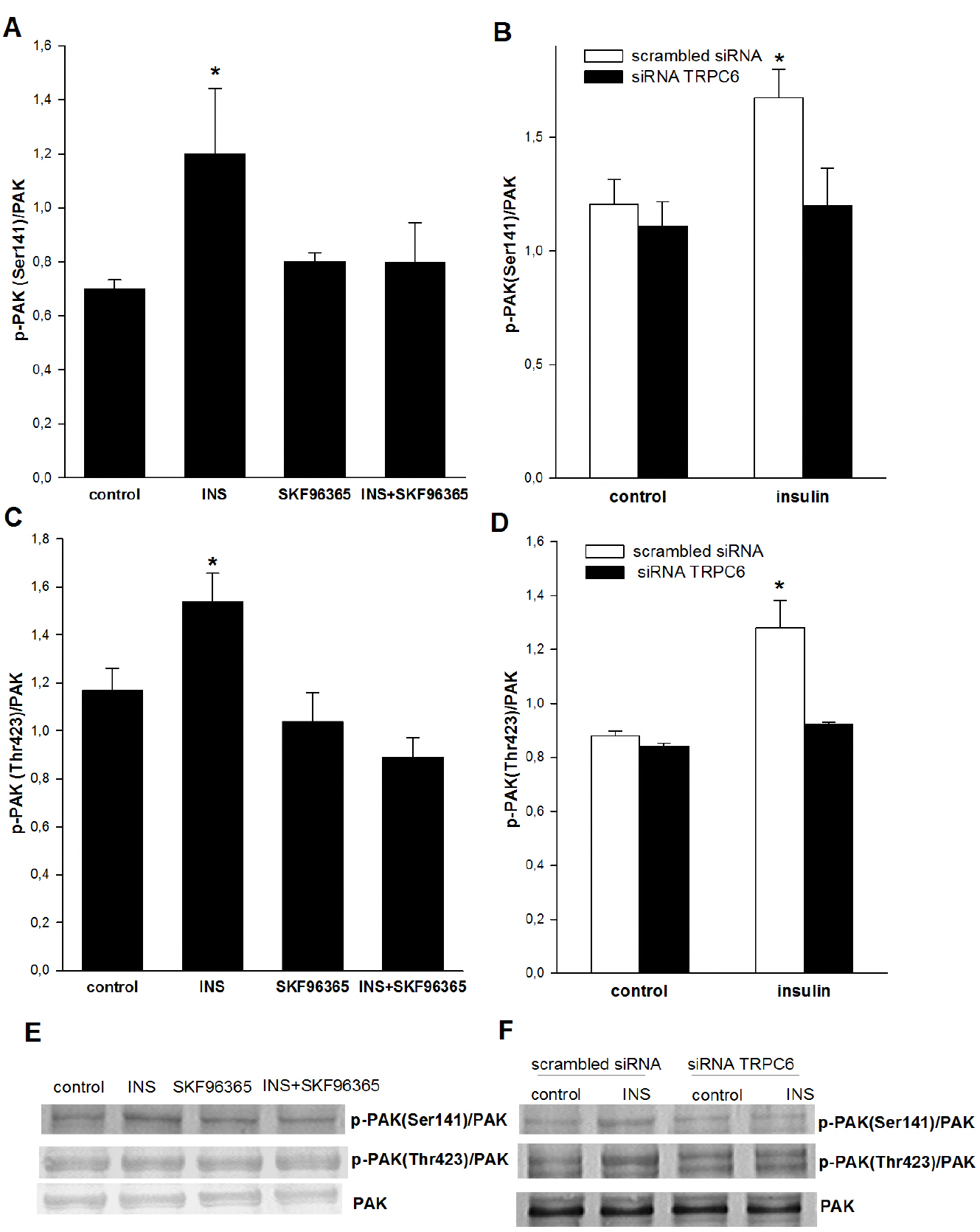
The role of TRPC channels in insulin-mediated phosphorylation of p21- activated kinase (PAK) in cultured rat podocytes. Podocytes were incubated for 5 min with 300 nM insulin in the presence or absence of the TRPC channel inhibitor SKF96365 (A, C) or TRPC6 siRNA (B, D). Densitometric analysis of the corresponding bands was performed, and values are reported as the ratios of the band intensities for p-PAK (Ser^141^) or p-PAK (Thr^423^) to PAK. Values are reported as the means±SEMs of four to six independent experiments. \**P*<0.05 compared to control.

**Figure 11.**
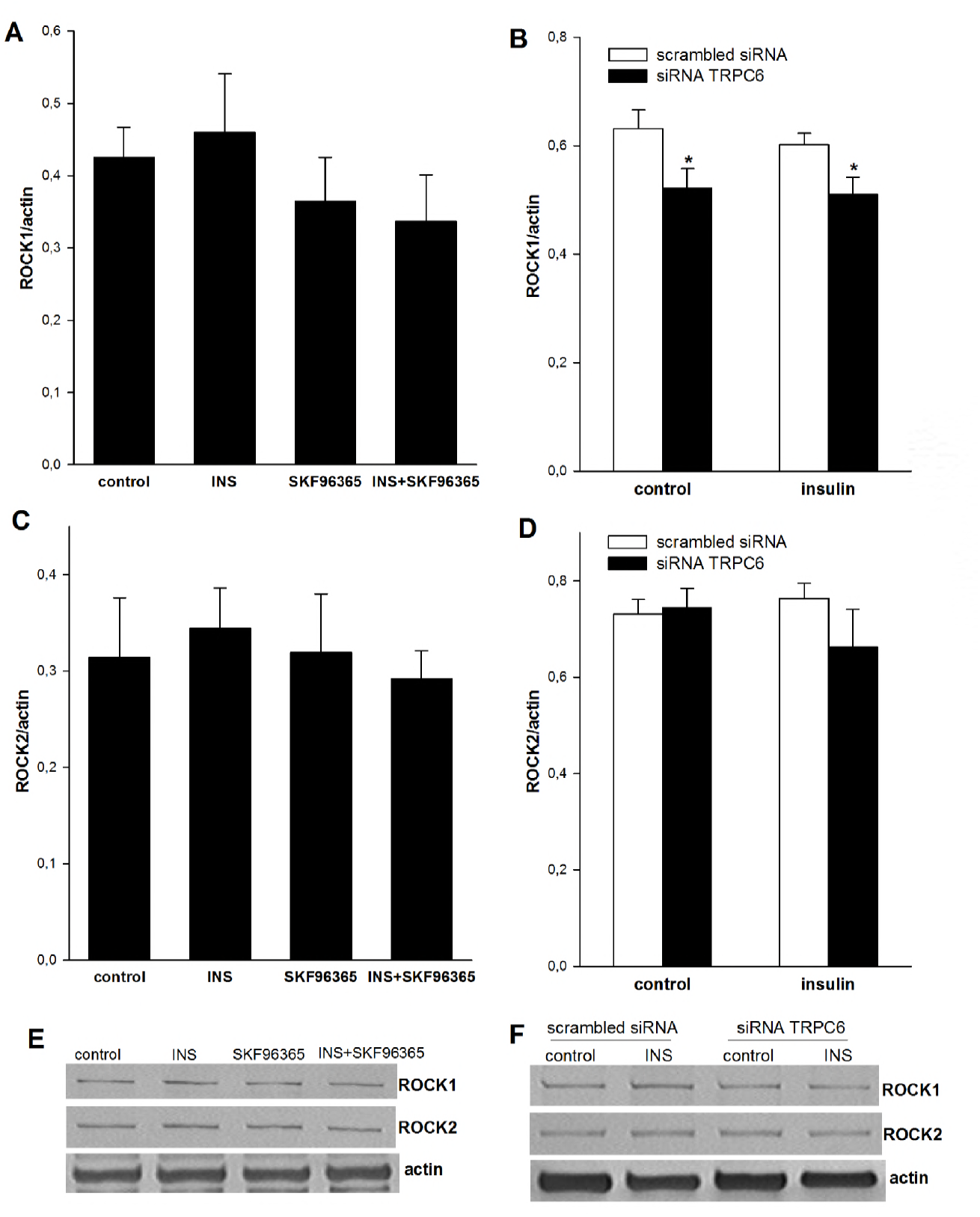
The influence of TRPC6 channels on ROCK1 and ROCK2 expression in podocytes. Podocytes were incubated for 5 min with 300 nM insulin in the presence or absence of the TRPC channel inhibitor SKF96365 (A, C) or TRPC6 siRNA (B, D). Densitometric analysis of the corresponding bands was performed, and values are reported as the ratios of the band intensities for ROCK1 or ROCK2 to actin. Values are reported as the means±SEMs of four to six independent experiments. \**P*<0.05 compared to control.

### TRPC6 channels regulate the insulin-dependent remodeling of the actin cytoskeleton in podocytes

The dynamics of actin filament assembly/disassembly and its organization in cells are regulated by several actin-binding proteins, including ADF/cofilins (Bamburg & Wiggan, 2002). In addition, PAK signals to cofilin in response to insulin, thereby facilitating cortical actin remodeling and glucose uptake in skeletal muscle cells (Tunduguru, Chiu et al., 2014). Accordingly, next we examined the effects of insulin on cofilin phosphorylation in podocytes. Notably, cofilin is activated when it is dephosphorylated. We found that insulin stimulation (300 nM, 5 min) decreased the p-cofilin level by about 25% (1.46±0.11 vs. control 1.95±0.10, *P*<0.05, Fig. 12) in cultured rat podocytes. Preincubation of the cells with SKF96365 (Fig. 12A) or downregulation of TRPC6 by siRNA (Fig. 12B) abolished the effect of insulin on cofilin phosphorylation.

**Figure 12.**
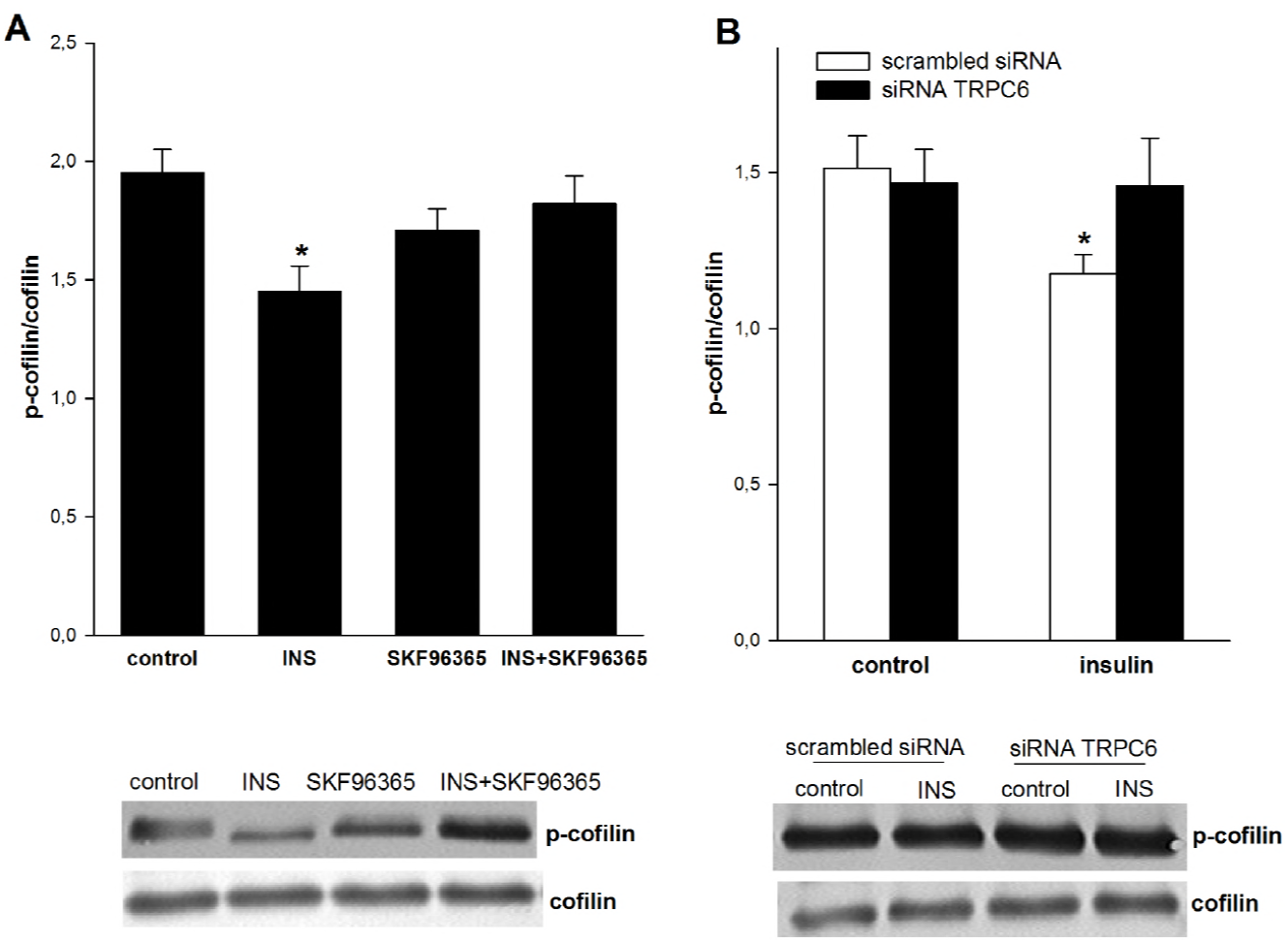
TRPC6 channel-mediated insulin-dependent activation of cofilin in cultured rat podocytes. The effects of the TRPC channel inhibitor SKF96365 (A) or TRPC6 siRNA (B) on the insulin-dependent dephosphorylation of cofilin. Values are reported as the means±SEMs of four to six independent experiments. \**P*<0.05 compared to untreated podocytes.

Phosphorylation of cofilin on serine3 leads to reduced actin binding and to actin depolymerization (Moriyama, Iida et al., 1996). We found that insulin increased the colocalization of cofilin with actin and that TRPC6 inhibition decreased this effect in podocytes (Fig. 13). Taken together, these results support accumulating evidence that TRPC6 participates in the regulation of actin dynamics in podocytes and suggest that this function is mediated in part by cofilin.

**Figure 13.**
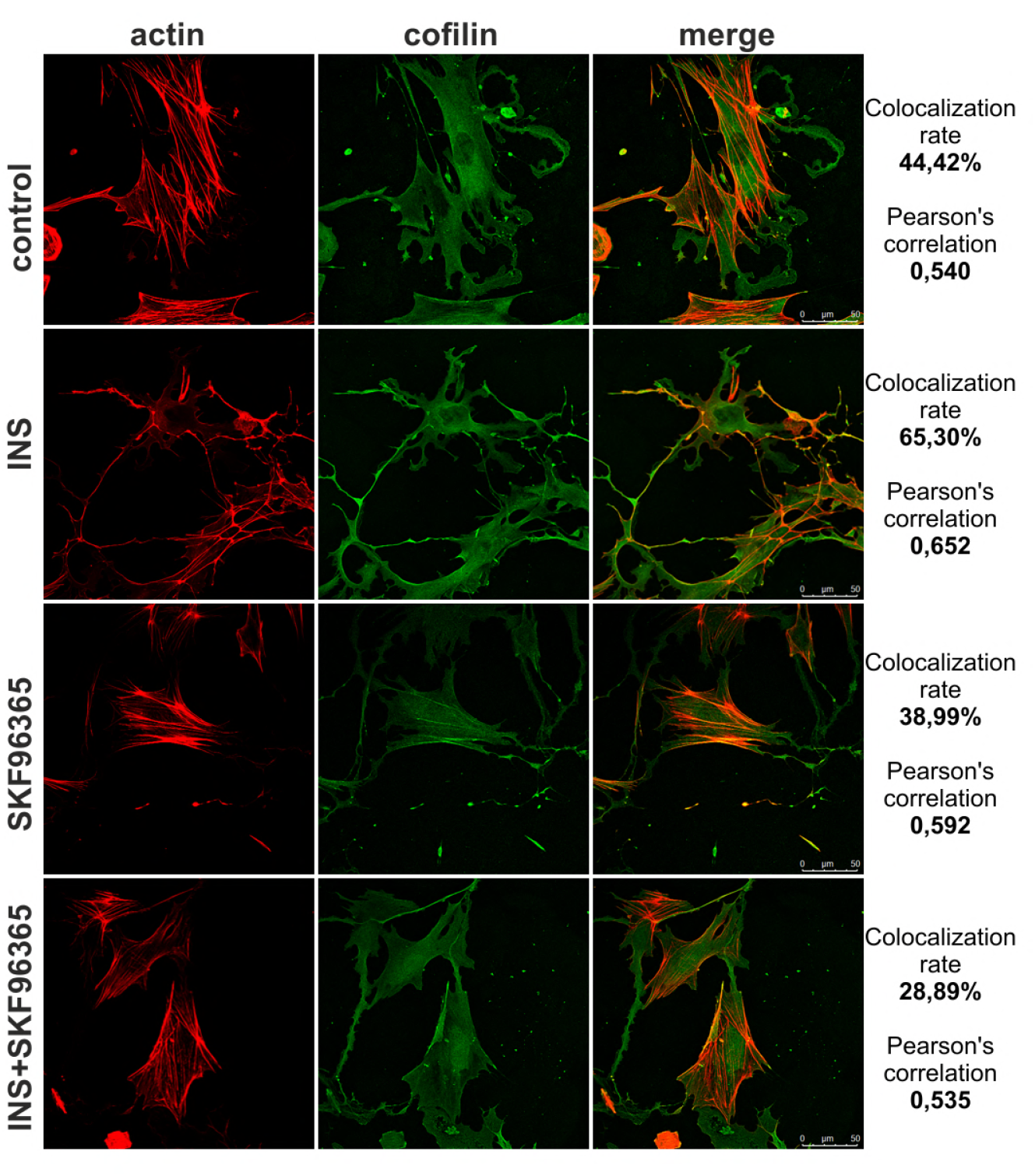
TRPC6 channels regulate insulin-induced changes in the colocalization of actin and cofilin. Rat podocytes seeded onto coverslips were incubated for 5 min with 300 nM insulin in the presence or absence of the TRPC channel inhibitor SKF96365. The cells were then immunostained with anti-cofilin antibody and isothiocyanate phalloidin to detect actin. Quantitative analysis of protein colocalization was performed with LAS AF 3.3.0 software (n=10). The pixel intensities were quantified, and the results are reported as Pearson’s correlation coefficients and colocalization rates (%).

## Discussion

This study showed that the TRPC6-dependent activation of AMPKα2 signaling pathways is a novel mechanism for the insulin-mediated regulation of actin cytoskeleton dynamics and glucose uptake in podocytes. The proposed mechanism, which is based on our data, is shown in Figure 14. First, insulin-stimulated glucose uptake and actin cytoskeleton reorganization depends on TRPC6 channel activation. Second, activation of the AMPKα signaling pathway is TRPC6-dependent. Third, insulin regulates the interaction of TRPC6 with AMPKα2 in cultured rat podocytes. Fourth, AMPK and TRPC6 activation are required to stimulate Rac1 signaling pathways.

**Figure 14.**
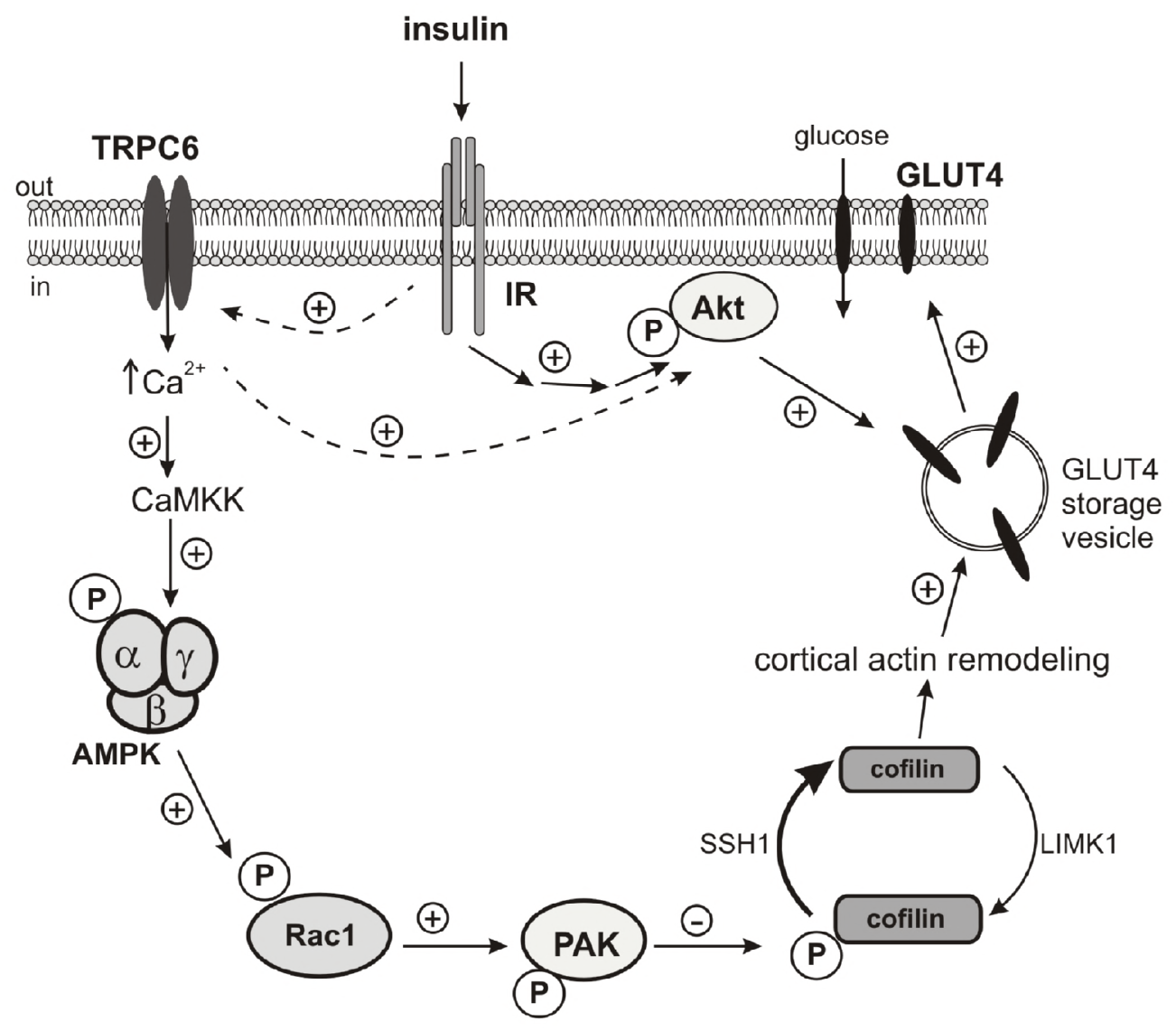
The proposed mechanism of insulin involvement in TRPC6-AMPK pathways that are involved in the cytoskeleton regulation and in the regulation of glucose uptake in podocytes.

We recently demonstrated that the insulin-mediated regulation of the contractile apparatus and filtration barrier permeability relies on TRPC6-dependent activation of PKGIα signaling pathways. Our studies found that insulin induced the reorganization of actin via TRPC6-dependent Ca^2+^ entry (Rogacka et al., 2017). The present study confirmed the role of TRPC6 in the regulation of insulin-dependent signaling pathways and actin dynamics. Moreover, we demonstrated for the first time that TRPC6 plays a role in the regulation of AMPK activity and glucose uptake in cultured rat podocytes.

AMPK is a major regulator of insulin-dependent glucose uptake and insulin signaling in podocytes (Rogacka et al., 2014). Phosphorylation of AMPK at Thr172 is required for the activation of AMPK (Hawley, Selbert et al., 1995), and Ca^2+^/calmodulin-dependent CaMKK-β activates AMPK in various cell types, including podocytes (Chen, Zhao et al., 2017, Nakanishi, Hatano et al., 2017, Piwkowska, Rogacka et al., 2011). CaMKK-β activity depends on increases in intracellular calcium, and insulin induces calcium flux into cells (Kim et al., 2012, Rogacka et al., 2017). Here we demonstrated that insulin induced a change in AMPK phosphorylation through TRPC6 activation. We also found, that insulin increased the amount of TRPC6 colocalized with the AMPKα2 subunit in podocytes. Others have also demonstrated a link between TRPC and AMPK activity. For example, TRPC1 knockdown in endothelial cells prevents PAR1 agonist peptide-induced AMPKα phosphorylation (Bair, Thippegowda et al., 2009). It is possible that AMPK and TRPC3 are part of the same signaling pathway that affects the cytoskeletal network and erythrocyte survival (Hirschler-Laszkiewicz, Tong et al., 2011). We assume that AMPK function depends on TRPC6-regulated calcium ion flux and we demonstrated that inhibiting TRPC6 channels and knocking down TRPC6 expression blocked insulin-dependent glucose uptake. We observed that the basal glucose uptake was not different between SKF96365-treated and TRPC6 siRNA-treated cells versus control cells; this is consistent with the fact that basal glucose uptake does not involve GLUT4 translocation.

Insulin and muscle contraction are the major known mediators of GLUT4 translocation under physiological conditions (Coward & Fornoni, 2015, Mueckler, 1994). The importance of podocyte insulin signaling in the pathogenesis of diabetic kidney disease is suggested by the observation that podocytes isolated from diabetic db/db mice cannot phosphorylate Akt in response to insulin and do not translocate GLUT4 to the plasma membrane after insulin stimulation (Guzman, Jauregui et al., 2014). We suggest that insulin induces the TRPC6-dependent entry of Ca^2+^ into podocytes, the translocation of GLUT4 to the membrane, and, consequently, an increase in glucose uptake in podocytes. However, this needs further investigation. Others have shown that TRPC3 knockdown reduces insulin-mediated glucose uptake in skeletal muscle cells. Moreover, TRPC3 and GLUT4 colocalize in the t-tubule system, which is responsible for insulin-dependent glucose uptake (Lanner, Bruton et al., 2009). It is clear that the ability of podocytes to precisely regulate intracellular Ca^2+^ levels plays a crucial role in proper glucose uptake.

The rho family GTPases act as molecular switches that are best known for their pivotal roles in the dynamic regulation of the actin cytoskeleton. The mammalian Rho family has at least 20 distinct members, of which RhoA, Rac1, and Cdc42 are the most extensively studied (Etienne-Manneville & Hall, 2002). Rac1 is a major regulator of actin remodeling, and cytoskeletal rearrangement is required for GLUT4 translocation in response to insulin (JeBailey et al., 2007).

These findings suggest that Rac1 and downstream signaling to the actin cytoskeleton constitute an important dysfunctional pathway in insulin-resistant states. In podocytes, cells that are sensitive to insulin, this mechanism may not be examined. The results presented here provide evidence that activation of AMPKα2 and TRPC6 are required for insulin-dependent Rac1 signaling pathway stimulation in podocytes. We further demonstrated that the insulin-mediated activation of downstream targets of Rho kinases was TRPC6 dependent. Inhibition of TRPC6 activity attenuated the effect of insulin on PAK phosphorylation, decreasing ROCK1 expression and cofilin activation. Insulin-stimulated Rac1 also activates the PAK1 serine/threonine kinase in skeletal muscle (Sylow et al., 2013). Moreover, PAK1 inhibition reduces insulin-induced cortical actin remodeling, cofilin activation, and GLUT4 translocation (Tunduguru et al., 2014). It is likely that the diminished activity of Rac1 and PAK1 in an insulin-resistant state leads to functional defects in proteins involved in actin remodeling that are necessary for glucose uptake in podocytes. Another group also demonstrated that the ability of insulin to increase glucose transport is attenuated by ROCK1 suppression (Chun, Araki et al., 2012). This suggests that a decrease in ROCK1 expression in podocytes after downregulation of TRPC6 additionally reduces insulin-dependent glucose transport.

Taken together, our data demonstrate that insulin induces the activation of PAK1 and cofilin-1, increases the colocalization of cofilin with actin, and, as a consequence, induces actin remodeling and glucose uptake. Cofilin disassembles actin, thereby maintaining the flexibility of actin remodeling and supporting the regeneration of free monomeric actin for further polymerization. This active cycling of actin as mediated by insulin is necessary for actin dynamics and to facilitate GLUT4 translocation to the surface of muscle cells [23].

In summary, our results add to accumulating evidence that TRPC6 participates in the insulin-mediated regulation of actin dynamics in podocytes and suggest that this function is mediated, at least in part, by cofilin. Thus, cofilin dephosphorylation might be attractive as a pharmacological target in order to ensure appropriate actin turnover in diabetes with insulin resistance and to ensure GLUT4 function and glucose uptake.

## Materials and Methods

### Preparation and culture of rat podocytes

All experimental procedures were performed in accordance with directive 2010/63/EU and were approved by the Local Bioethics Commission at the Medical University of Gdansk. We used female Wistar rats weighting 100–120 g. Podocytes were isolated as described previously (Piwkowska, Rogacka et al., 2012). Experiments were conducted using podocytes cultivated for 12–20 days. Cell phenotypes were established using podocyte-specific antibodies against Wilms tumor-1 protein (WT-1; Biotrend Koeln, Germany) and synaptopodin (Progen, Heidelberg, Germany).

### Western blot analysis

To obtain podocyte lysate, the cells were treated with lysis buffer (1% Nonidet P-40, 20 mM Tris, 140 mM NaCl, 2 mM EDTA, 10% glycerol) in the presence of a protease inhibitor cocktail and homogenized at 4°C by scraping. Proteins in the supernatant (12 μg) were separated on a 10% SDS-polyacrylamide gel and electrotransferred to nitrocellulose membranes. The following primary antibodies were used for Western blotting: anti-p-AMPKα (Thr172) (1:1000, Cell Signaling Technology), anti-AMPKα (1:1000, Cell Signaling Technology), anti-p-insulin Rβ (Tyr1150-1151) (1:200, Santa Cruz Biotechnology), anti-insulin Rβ (1:200, Santa Cruz Biotechnology), anti-p-Akt1/2/3 (Ser473) (1:400, Santa Cruz Biotechnology), anti-Akt1/2/3 (1:400, Santa Cruz Biotechnology), anti-TRPC6 (1:1000, Sigma-Aldrich), anti-p-PAK 1/2/3 (Thr 423/402/421) (1:800, Sigma-Aldrich), anti-p-PAK1/2/3 (Ser144/141/139), anti-PAK1/2/3 (1:800, Cell Signaling Technology), anti-ROCK1 (1:1000, Cell Signaling Technology), anti-ROCK2 (1:1000, Cell Signaling Technology), anti-p-cofilin (1:1000, Sigma-Aldrich), anti-cofilin (1:1000, Santa Cruz Biotechnology), p-Rac1 (Ser71) (1:1000, OriGene), and anti-actin (1:3000, Sigma-Aldrich). To detect the primary antibodies, the membranes were incubated with the appropriate alkaline phosphatase-labeled secondary antibodies. The protein bands were visualized using the colorimetric 5-bromo-4-chloro-3-indolylphasphate/nitroblue tetrazolium (BCIP/NBT) system.

### Measurement of glucose uptake

Glucose uptake was measured as described previously (Rogacka et al., 2016) by the addition of 1 μCi/well of (1,2-3H)-deoxy-D-glucose diluted in non-radioactive glucose to a final concentration of 50 μM with or without 300 nM insulin. After incubation for 3 min, intracellular and extracellular radioactivity was measured by liquid scintillation counting using a MicroBeta2 Microplate Counter (Perkin Elmer, Waltham, MA, USA).

### RNA interference and cell transfection

Podocytes were transfected with small interfering RNAs (siRNAs) that targeted TRPC6 (OriGene), AMPKα1, AMPKα2 (Santa Cruz Biotechnology) or with control, non-silencing siRNA (scrambled siRNA, negative control) (OriGene or Santa Cruz Biotechnology). Cells were cultured in RPMI 1640 supplemented with 10% FBS. One day before the experiment, the culture medium was changed to antibiotic-free RPMI 1640 supplemented with 10% FBS. The cells were transfected with siRNAs using siRNA Transfection Reagent (Santa Cruz Biotechnology) according to the manufacturer’s instructions. Briefly, the targeted siRNA or scrambled siRNA were diluted in Transfection Medium (final concentration, 80 nM), mixed with siRNA Transfection Reagent, and incubated for 30 min at room temperature. Then Transfection Medium was added to the transfection mixture, mixed gently, and added to the podocytes. After 7 h, we added grow medium supplemented with 2× higher concentrations of FBS and antibiotics. The podocytes were incubated for an additional 24 h. After transfection, gene silencing was checked at the protein level by Western blotting.

### Immunofluorescence

Podocytes were seeded on coverslips coated with type 1 collagen (Becton Dickinson Labware, Becton, UK) and cultured in RPMI 1640 supplemented with 10% FBS. Cells were fixed in PBS plus 4% formaldehyde for 10 min at room temperature. Fixed podocytes were permeabilized with 0.3% Triton-X for 3–4 min and then blocked with PBSB solution (PBS plus 2% FBS, 2% BSA, and 0.2% fish gelatin) for 1 h. After blocking, cells were incubated with anti-cofilin (1:50), anti-TRPC6 (1:100), anti-AMPKα1 (1:100), and anti-AMPKα2 (1:100) antibodies in PBSB at 4°C for 1 h. The primary antibodies were incubated with blocking peptide to eliminate non-specific staining. Next, the cells were washed three times with cold PBS and incubated with secondary antibodies conjugated to Alexa Fluor 488 (1:1000) or Alexa Fluor 546 (1:1000). Actin was stained using Alexa Fluor 594 phalloidin (1:200). Specimens were imaged using a confocal laser scanning microscope (Leica SP8X) with a 63× oil immersion lens.

### Immunoprecipitation

Cell extracts were pre-cleared with mouse IgG plus Protein A/G-PLUS Agarose at 4°C for 1 h and then incubated with a primary antibody plus Protein A/G-PLUS Agarose at 4°C overnight. The agarose beads were washed gently with lysis buffer. Proteins were eluted from the beads by adding SDS loading buffer. The mixture was then boiled for 5 min and subjected to Western blot analysis.

### Rac1 activity assay

Rac1 activation was measurement in the supernatant using a commercially available G-LISA Rac1 Activation Assay Biochem Kit (BK128; Cytoskeleton, Inc.)

### Statistical analysis

Statistical analyses were performed using one-way ANOVA followed by the Student-Newman-Keuls test to determine significance. Values are reported as means ± SEMs. Significance was set at *P*<0.05.

## Acknowledgments

This work was supported by grants from the National Science Center (grants 2014/14/E/NZ4/00358 and 2015/17/B/NZ4/02658).

## Author contribution

Conceptualisation: SA and AP; Investigation: PR, MSz, IA, DR, MR and AP; writing—original draft: PR and AP; writing—review and editing: DR, IA, SA and AP; visualization: PR and MR; supervision: SA and AP.

## References

Bair AM, Thippegowda PB, Freichel M, Cheng N, Ye RD, Vogel SM, Yu Y, Flockerzi V, Malik AB, Tiruppathi C (2009) Ca2+ entry via TRPC channels is necessary for thrombin-induced NF-kappaB activation in endothelial cells through AMP-activated protein kinase and protein kinase Cdelta. J Biol Chem 284: 563–74

Bamburg JR, Wiggan OP (2002) ADF/cofilin and actin dynamics in disease. Trends Cell Biol 12: 598–605

Burridge K, Wennerberg K (2004) Rho and Rac take center stage. Cell 116: 167–79

Chen X, Zhao X, Lan F, Zhou T, Cai H, Sun H, Kong W (2017) Hydrogen Sulphide Treatment Increases Insulin Sensitivity and Improves Oxidant Metabolism through the CaMKKbeta-AMPK Pathway in PA-Induced IR C2C12 Cells. Sci Rep 7: 13248

Chiu TT, Patel N, Shaw AE, Bamburg JR, Klip A (2010) Arp2/3- and cofilin-coordinated actin dynamics is required for insulin-mediated GLUT4 translocation to the surface of muscle cells. Mol Biol Cell 21: 3529–39

Chun KH, Araki K, Jee Y, Lee DH, Oh BC, Huang H, Park KS, Lee SW, Zabolotny JM, Kim YB (2012) Regulation of glucose transport by ROCK1 differs from that of ROCK2 and is controlled by actin polymerization. Endocrinology 153: 1649–62

Coward R, Fornoni A (2015) Insulin signaling: implications for podocyte biology in diabetic kidney disease. Curr Opin Nephrol Hypertens 24: 104–10

Coward RJ, Welsh GI, Yang J, Tasman C, Lennon R, Koziell A, Satchell S, Holman GD, Kerjaschki D, Tavare JM, Mathieson PW, Saleem MA (2005) The human glomerular podocyte is a novel target for insulin action. Diabetes 54: 3095–102

Dryer SE, Reiser J (2010) TRPC6 channels and their binding partners in podocytes: role in glomerular filtration and pathophysiology. Am J Physiol Renal Physiol 299: F689–701

Etienne-Manneville S, Hall A (2002) Rho GTPases in cell biology. Nature 420: 629–35

Fisher JS, Gao J, Han DH, Holloszy JO, Nolte LA (2002) Activation of AMP kinase enhances sensitivity of muscle glucose transport to insulin. Am J Physiol Endocrinol Metab 282: E18–23

Guzman J, Jauregui AN, Merscher-Gomez S, Maiguel D, Muresan C, Mitrofanova A, Diez-Sampedro A, Szust J, Yoo TH, Villarreal R, Pedigo C, Molano RD, Johnson K, Kahn B, Hartleben B, Huber TB, Saha J, Burke GW, 3rd, Abel ED, Brosius FC et al. (2014) Podocyte-specific GLUT4-deficient mice have fewer and larger podocytes and are protected from diabetic nephropathy. Diabetes 63: 701–14

Hawley SA, Selbert MA, Goldstein EG, Edelman AM, Carling D, Hardie DG (1995) 5’-AMP activates the AMP-activated protein kinase cascade, and Ca2+/calmodulin activates the calmodulin-dependent protein kinase I cascade, via three independent mechanisms. J Biol Chem 270: 27186–91

Hirschler-Laszkiewicz I, Tong Q, Waybill K, Conrad K, Keefer K, Zhang W, Chen SJ, Cheung JY, Miller BA (2011) The transient receptor potential (TRP) channel TRPC3 TRP domain and AMP-activated protein kinase binding site are required for TRPC3 activation by erythropoietin. J Biol Chem 286: 30636–46

JeBailey L, Wanono O, Niu W, Roessler J, Rudich A, Klip A (2007) Ceramide-and oxidant-induced insulin resistance involve loss of insulin-dependent Rac-activation and actin remodeling in muscle cells. Diabetes 56: 394–403

Jiang L, Ding J, Tsai H, Li L, Feng Q, Miao J, Fan Q (2011) Over-expressing transient receptor potential cation channel 6 in podocytes induces cytoskeleton rearrangement through increases of intracellular Ca2+ and RhoA activation. Exp Biol Med (Maywood) 236: 184–93

Jing M, Cheruvu VK, Ismail-Beigi F (2008) Stimulation of glucose transport in response to activation of distinct AMPK signaling pathways. Am J Physiol Cell Physiol 295: C1071–82

Kim EY, Anderson M, Dryer SE (2012) Insulin increases surface expression of TRPC6 channels in podocytes: role of NADPH oxidases and reactive oxygen species. Am J Physiol Renal Physiol 302: F298–307

Kim EY, Dryer SE (2011) Effects of insulin and high glucose on mobilization of slo1 BKCa channels in podocytes. J Cell Physiol 226: 2307–2308

Krall P, Canales CP, Kairath P, Carmona-Mora P, Molina J, Carpio JD, Ruiz P, Mezzano SA, Li J, Wei C, Reiser J, Young JI, Walz K (2010) Podocyte-specific overexpression of wild type or mutant trpc6 in mice is sufficient to cause glomerular disease. PLoS One 5: e12859

Lanner JT, Bruton JD, Assefaw-Redda Y, Andronache Z, Zhang SJ, Severa D, Zhang ZB, Melzer W, Zhang SL, Katz A, Westerblad H (2009) Knockdown of TRPC3 with siRNA coupled to carbon nanotubes results in decreased insulin-mediated glucose uptake in adult skeletal muscle cells. Faseb j 23: 1728–38

Lei M, Lu W, Meng W, Parrini MC, Eck MJ, Mayer BJ, Harrison SC (2000) Structure of PAK1 in an autoinhibited conformation reveals a multistage activation switch. Cell 102: 387–97

Mathieson PW (2011) The podocyte as a target for therapies-new and old. Nat Rev Nephrol 8: 52–6

Moriyama K, Iida K, Yahara I (1996) Phosphorylation of Ser-3 of cofilin regulates its essential function on actin. Genes Cells 1: 73–86

Mueckler M (1994) Facilitative glucose transporters. Eur J Biochem 219: 713–25

Nakanishi A, Hatano N, Fujiwara Y, Sha’ri A, Takabatake S, Akano H, Kanayama N, Magari M, Nozaki N, Tokumitsu H (2017) AMP-activated protein kinase-mediated feedback phosphorylation controls the Ca(2+)/calmodulin (CaM) dependence of Ca(2+)/CaM-dependent protein kinase kinase beta. J Biol Chem 292: 19804–19813

Piwkowska A, Rogacka D, Audzeyenka I, Kasztan M, Angielski S, Jankowski M (2015) Insulin increases glomerular filtration barrier permeability through PKGIalpha-dependent mobilization of BKCa channels in cultured rat podocytes. Biochim Biophys Acta 1852: 1599–609

Piwkowska A, Rogacka D, Audzeyenka I, Kasztan M, Angielski S, Jankowski M (2016) Intracellular calcium signaling regulates glomerular filtration barrier permeability: the role of the PKGIα- dependent pathway. FEBS Lett 590: 1739–48

Piwkowska A, Rogacka D, Jankowski M, Angielski S (2011) Extracellular ATP through P2 receptors activates AMP-activated protein kinase and suppresses superoxide generation in cultured mouse podocytes. Exp Cell Res 317: 1904–13

Piwkowska A, Rogacka D, Jankowski M, Kocbuch K, Angielski S (2012) Hydrogen peroxide induces dimerization of protein kinase G type Ialpha subunits and increases albumin permeability in cultured rat podocytes. J Cell Physiol 227: 1004–16

Piwkowska A, Rogacka D, Kasztan M, Angielski S, Jankowski M (2013) Insulin increases glomerular filtration barrier permeability through dimerization of protein kinase G type Ialpha subunits. Biochim Biophys Acta 1832: 791–804

Rawshani A, Gudbjornsdottir S (2017) Mortality and Cardiovascular Disease in Type 1 and Type 2 Diabetes. In N Engl J Med, pp 300–301. United States:

Ritz E, Rychlik I, Locatelli F, Halimi S (1999) End-stage renal failure in type 2 diabetes: A medical catastrophe of worldwide dimensions. Am J Kidney Dis 34: 795–808

Rogacka D, Audzeyenka I, Rachubik P, Rychlowski M, Kasztan M, Jankowski M, Angielski S, Piwkowska A (2017) Insulin increases filtration barrier permeability via TRPC6-dependent activation of PKGIalpha signaling pathways. Biochim Biophys Acta 1863: 1312–1325

Rogacka D, Piwkowska A, Audzeyenka I, Angielski S, Jankowski M (2014) Involvement of the AMPK-PTEN pathway in insulin resistance induced by high glucose in cultured rat podocytes. Int J Biochem Cell Biol 51: 120–30

Rogacka D, Piwkowska A, Audzeyenka I, Angielski S, Jankowski M (2016) SIRT1-AMPK crosstalk is involved in high glucose-dependent impairment of insulin responsiveness in primary rat podocytes. Exp Cell Res 349: 328–338

Sylow L, Jensen TE, Kleinert M, Hojlund K, Kiens B, Wojtaszewski J, Prats C, Schjerling P, Richter EA (2013) Rac1 signaling is required for insulin-stimulated glucose uptake and is dysregulated in insulin-resistant murine and human skeletal muscle. Diabetes 62: 1865–75

Sylow L, Moller LL, Kleinert M, Richter EA, Jensen TE (2015) Stretch-stimulated glucose transport in skeletal muscle is regulated by Rac1. J Physiol 593: 645–56

Sylow L, Moller LLV, Kleinert M, D’Hulst G, De Groote E, Schjerling P, Steinberg GR, Jensen TE, Richter EA (2017) Rac1 and AMPK Account for the Majority of Muscle Glucose Uptake Stimulated by Ex Vivo Contraction but Not In Vivo Exercise. Diabetes 66: 1548–1559

Tsakiridis T, Taha C, Grinstein S, Klip A (1996) Insulin activates a p21-activated kinase in muscle cells via phosphatidylinositol 3-kinase. J Biol Chem 271: 19664–7

Tunduguru R, Chiu TT, Ramalingam L, Elmendorf JS, Klip A, Thurmond DC (2014) Signaling of the p21- activated kinase (PAK1) coordinates insulin-stimulated actin remodeling and glucose uptake in skeletal muscle cells. Biochem Pharmacol 92: 380–8

Welsh GI, Hale LJ, Eremina V, Jeansson M, Maezawa Y, Lennon R, Pons DA, Owen RJ, Satchell SC, Miles MJ, Caunt CJ, McArdle CA, Pavenstadt H, Tavare JM, Herzenberg AM, Kahn CR, Mathieson PW, Quaggin SE, Saleem MA, Coward RJ (2010) Insulin signaling to the glomerular podocyte is critical for normal kidney function. Cell Metab 12: 329–40

Zeqiraj E, Filippi BM, Deak M, Alessi DR, van Aalten DM (2009) Structure of the LKB1-STRAD-MO25 complex reveals an allosteric mechanism of kinase activation. Science 326: 1707–11

